# Temporal changes in the protein cargo of extracellular vesicles and resultant immune reprogramming after severe burn injury in humans and mice

**DOI:** 10.1101/2025.03.19.644202

**Authors:** Micah L. Willis, Roland Seim, Laura E. Herring, Angie L. Mordant, Thomas S. Webb, Gilbert R. Upchurch, Ashish K. Sharma, Bruce A. Cairns, Philip Efron, Shannon M. Wallet, Leon G. Coleman, Robert Maile

## Abstract

**Introduction:** Severe injury, including burn trauma, leads to profound immune dysfunction, yet the mechanisms driving these changes remain incompletely defined. This lack of understanding has hindered efforts to modulate the immune response effectively. Additionally, a clear biomarker profile to guide clinicians in identifying burn patients at high risk for poor clinical outcomes is lacking. Extracellular vesicles (EVs) have emerged as novel mediators of immune dysfunction in various pathologies. Prior studies in mouse models have demonstrated that plasma EVs increase following burn injury and contribute to immune dysfunction. Furthermore, EVs have potential as biomarkers for predicting extended hospital stays in burn patients. This study hypothesizes that human EVs, purified early and late after burn injury, will exhibit immune reprogramming effects similar to those observed in mice and that specific EV protein cargo may serve as biomarkers of immune and physiological responses to burn injury.

**Methods:** EVs were isolated from the plasma of burn-injury patients at early (<72h) and late (≥14 days) time points post-injury. Using unbiased immune transcriptome and bioinformatic causal network analyses, the immunomodulatory effects of these EVs were assessed in human THP-1 macrophages. Mass spectrometry-based quantitative proteomics and pathway analyses were conducted to characterize the protein cargo of EVs from both human and mouse models at different post-burn phases.

**Results:** Early post-burn human EVs induced significant immune reprogramming in macrophages, increasing pro-inflammatory signaling while suppressing anti-inflammatory pathways. In contrast, late post-burn EVs exhibited an immunosuppressive profile, with downregulation of pro-inflammatory pathways and upregulation of anti-inflammatory signaling. Proteomic analyses revealed that human and mouse EVs contained unique and overlapping protein cargo across different time points. At day 7 post-burn, mouse EVs were enriched in circulation/complement and neuronal proteins, whereas by day 14, reductions in membrane and metabolism-associated proteins were observed. Similarly, in human EVs at 14 days post-burn, increased levels of circulation/complement, immune, and transport proteins were detected.

**Conclusions:** EVs from burn-injury patients at distinct time points differentially modulate immune responses in macrophages, mirroring the temporal immune phenotypes observed in clinical settings. These findings suggest that EV-macrophage interactions play a crucial role in burn-induced immune dysfunction and highlight the potential of EV protein cargo as biomarkers for immune status and patient outcomes following burn injury.

**Summary Sentence:** Human extracellular vesicles released into the plasma after severe burn injury can reprogram the immune system with corresponding immunomodulatory protein cargo.

## INTRODUCTION

Severe burn injury presents itself as biphasic system of immune dysfunction, with an acute phase (0-72 hours post injury) comprising a ‘Systemic Inflammatory Response Syndrome’ (SIRS; “burn shock”) with concurrent coagulopathy, metabolic dysfunction, and multiple organ failure (1). Following this phase, a period of time occurs associated with profound susceptibility to hospital-acquired infections (HAI)(2). This clinically important period is defined as a Compensatory Anti-Inflammatory Response Syndrome (CARS) but over time has been better defined as a Persistent Inflammation, Immunosuppression, and Catabolism Syndrome (PICS). The initiating and controlling mechanisms for SIRS and PICS have not been fully elucidated and attempts to modulate these responses have been generally unsuccessful (for example, (3)). Additionally, there is a lack of a clear biomarker profile that would guide clinicians in the care of burn patients to identify patients at high risk for poor clinical outcomes.

Recently, extracellular vesicles (EVs) have emerged as novel mediators of immune dysfunction across several immune pathologies, including our own work in burn-injury, reviewed in *Shock* (4). EVs are released from nearly all cell types with their size, concentration, and composition based on pathological conditions. EVs are classified primarily based on their size and biogenesis into three main categories: apoptotic bodies (>1µm), microvesicles (∼0.1-1µm), and exosomes (∼50-100 nm) (4, 5). There is some overlap in size, cargo, and surface markers between exosomes and EVs, though exosomes originate from the endosomal system whereas EVs primarily arise from budding from cell surface membranes (4, 5). Previously, we have reported using a mouse model of burn injury and burned patients that a mixture of altered exosomes and microvesicles arise after burn injury ((6, 7)). Using the mouse model, we demonstrated clear immunomodulatory effects on immune cells by EVs harvested early (1 day) and late after injury (14 days). In both cases we characterized protein cargo by unbiased mass spectrometry in EVs that arose early after injury. We found cargo overlap exist between humans and mice which demonstrated their potential as biomarkers in burn patients; early after burn injury EV were enriched with both serum amyloid A1 (SAA1) and C-reactive protein (CRP) which correlated with eventual length of hospital stay ((6)).

We have reported that early after burn injury (<72h) in human patients and mice that there is a higher frequency of EVs loaded with immunomodulators, such as HMGB1 and IL-1ß, compared to EVs isolated from healthy humans or sham injured mice (8). We have also demonstrated that EV from the burned mouse harvested early and late after injury were able to induce significant alterations in immune gene expression in macrophages (6, 7). Differential immune phenotypes were observed depending on the time after injury that the EV were harvested (6, 7). To further determine EV signatures that arise after burn injury, we have shown that plasma EVs released <u>late</u> after burn injury continue to promote immune dysfunction. Further protein cargo analysis of EV that arise late after burn injury revealed specific immune and physiologic pathways that these protein cargo impact. In both cases, data highlighted that EV purified late after burn injury differ to EV purified early after injury in that they are capable of promoting immune pathways associated with the profound immune susceptibility to infection experienced by burn patients and mice later after injury.

## METHODS

### Mouse 20% TBSA Thermal Injury

All procedures were conducted in strict adherence to the Guide for the Care and Use of Laboratory Animals of the National Institute of Health and approved by the University of North Carolina Institutional Animal Care and Use Committee under protocol #21-082. Mice were housed in American Association for Accreditation of Laboratory Animal Care (AAALAC)-accredited facilities with 24 h veterinary care and close observation throughout the experiment. Measures were taken to alleviate suffering, with all injuries performed under avertin anesthesia with morphine post-burn analgesia. Mice underwent a 20% total body surface area (TBSA) thermal injury to model a large burn injury in humans as described previously (7). Briefly, C57BL6 mice (female, 6–8 weeks old, 15–20 g) first were anesthetized with tribromoethanol/avertin (475 mg/kg; Sigma-Aldrich, Burlington MA, USA). Their targeted region on the dorsum was then shaved (NC0854145; Fisher, Pittsburg, PA, USA) prior to injection of subcutaneous morphine sulfate (3 mg/kg; Westward, Berkeley Heights, NJ, USA). Four defined skin locations were then contacted for 10 s with a copper rod heated to 100 °C in a water bath. Mice were then resuscitated with Ringer’s lactate (0.1 mL/g body weight). Post-procedure analgesia was maintained with morphine sulfate-supplemented water (60 µg/20 g mouse) ad lib for the duration of the experiment. Sham mice underwent identical treatment minus application of a heated copper rod. Mice were monitored at least twice daily for the duration of the experiment. Mice that showed signs of distress including a >15% loss of body weight, difficulty breathing, hunching over, dehydration, inactivity, or growing lesions were humanely euthanized immediately. There was zero burn-related mortality.

### EV Isolation, Quantification, and Sizing

EVs were isolated from plasma and centrifuged at 2,000 x g for 20 minutes to remove cells. Supernatant was then centrifuged at 10,000 x g for 30 minutes to remove cellular debris. Remaining supernatant was then centrifuged at 21,000 x g for 1 hour. The EV-containing pellet was washed in PBS and centrifuged again at 21,000 x g. This preparation results in EVs ranging between 100 nm and 1 μm in diameter (9). EV pellets were resuspended in 2-3 mL of saline, filtered with a 0.22 uM syringe filter, and frozen at –80°C. Nanoparticle Tracking Analysis (NTA) was performed on the final EV products using the ZetaView QUATT instrument (Particle Metrix) and ZetaView (version 8.05) software. Mean concentrations (EV/mL) and mode size (diameter in nanometers) were determined from 10 videos taken of one sample analyzed at a 1:1000 dilution with filtered PBS with a 488nm laser, pump speed 30, camera shutter of 100. Each measurement obtained from the 10 videos were internally quality controlled by the instrument, with videos removed for failing quality control.

### Macrophage Culture and EV Stimulation

THP-1 (ATCC, Manassas, VA, USA) human monocytic cells were allowed to grow in culture according to manufacturer’s instructions using RPMI 1640 media containing 10% FBS, 0.05 mM 2-mercaptoethanol, and 1% penicillin/streptomycin at 37°C and 5% CO_2_. A total of 5 x 10^5^ cells were plated in a 24-well cell culture plates and stimulated with 200 nM phorbol 12-myristate-13-acetate (PMA) for 24 hours. The cell media was removed and replaced with fresh media and allowed to rest for 24 hours. Resultant cells, which are predominantly conditioned into macrophages, were exposed to 3 x 10^7^ EVs in the absence or presence of LPS from *Escherichia coli* O111:B4 for 48 hours. Supernatants and cellular mRNA were harvested for analysis.

### Subject Characteristics

Blood samples from burn patients admitted to the North Carolina Jaycee Burn Center and recruited into an IRB-approved repository protocol (IRB 04-1437) were collected and stored. Patients received standard of care and care was not affected by study participation. Patients were not excluded from the study based on burn size, inhalation injury or factors including age, race, or substance use prior to injury. Patients were followed until discharge or expiration. This study utilized plasma samples collected early after injury (1-3 days) and late after injury (18-25 days) from recruited burn patients (Table 1).

**Table 1.** Clinical features of burn patients, n=15 (LOS=Length of Stay).

| % TBSA Injury | Age | Sex % | LOS | Mortality # (%) |
| --- | --- | --- | --- | --- |
| 18.4 ± 1.7,<br>n=15 | 50.3 ± 2.2 | M- 68<br>F – 32 | 49.4 ± 8.9 | Overall – 4 (8)<br>M – 2 (5.2)<br>F- 2 (12.5) |

### Human chemokine and cytokine analysis

Bio-Plex Human Screening Panel 8-Plex (BioRad 17009195) was used to probe cell supernatants for IL-1β, IL-2, IL-6, IL-10, IL-12(p70), IFN-ᵧ, CCL2 (MCP-1) and TNF according to manufacturer’s protocol. Data was acquired on a Bio-Plex MAGPIX Multiplex reader system (BioRad Hercules, California, USA) running xPONENT and analyzed using a using a parameter logistic spine-curve fitting method. All data are presented as picograms per milliliter.

### Immune gene expression detection and quantification

Isolation of mRNA was performed as previously (8). Briefly, THP-1 macrophages were lysed with TRIZOL buffer (Sigma) and total RNA was isolated by chloroform extraction and quantified using a nanodrop 2000TM(Wilmington DE) spectrophotometer. NanoString technology and the nCounter Human Immunology Panel v2 was used to simultaneous evaluate 561 mRNAs in each sample (10). Each sample was run in triplicate. nSolver v4.0, an integrated analysis platform was used to generate appropriate data normalization as well as fold-changes, resulting ratios and differential expression. nCounter™ v4.0 Advanced Analysis and Ingenuity Pathway Analysis along with R were used to identify pathway-specific responses (10).

### Unbiased proteomic assessment of EVs from mice and humans and using LC-MS/MS

EVs were isolated from sham injured mice, 20% TBSA burn injury mice, healthy humans, and human burn patients. Plasma from mice was collected 1, 7 and 14 days after injury, and plasma from human burn patients within the first 72 hours and 18-25 days of injury/hospital admission and following discharge. After the final spin, EVs were resuspended in 20 mM Tris buffer (pH 7.5). The UNC Proteomics Core’s proteomics workflow was applied as described previously (11–13). Briefly, 8 M urea was added to the in-solution protein samples (∼10-20 µg per replicate, n = 3), then reduced with 5 mM DTT for 30 min at 37ᵒ C and alkylated with 15 mM iodoacetamide for 45 min in the dark at room temperature. The samples were diluted to 1 M urea, then digested with MS grade trypsin (Promega) at 37°C overnight. The peptide samples were acidified to 1% TFA, then desalted using C18 desalting spin columns (Pierce). Peptide concentration was determined using a fluorometric peptide quantitation assay (Pierce). Samples were dried via vacuum centrifugation and reconstituted, normalizing to 0.1 ug/μL. LC-MS/MS Analysis: Each sample was analyzed by LC-MS/MS using an Easy nLC 1200 coupled to a QExactive HF (Thermo Scientific). Samples were injected onto an Easy Spray PepMap C18 column (75 μm id × 25 cm, 2 μm particle size) (Thermo Scientific) and separated over a 90 min method. The gradient for separation consisted of 5–32% mobile phase B at a 250 nl/min flow rate, where mobile phase A was 0.1% formic acid in water and mobile phase B consisted of 0.1% formic acid in ACN. The QExactive HF was operated in data-dependent mode where the 15 most intense precursors were selected for subsequent HCD fragmentation. Resolution for the precursor scan (m/z 350–1700) was set to 60,000 with a target value of 3 × 106 ions, 100 ms inject time. MS/MS scans resolution was set to 15,000 with a target value of 1 × 105 ions, 75 ms inject time. The normalized collision energy was set to 27% for HCD, with an isolation window of 1.6 m/z. Peptide match was set to preferred, and precursors with unknown charge or a charge state of 1 and ≥ 7 were excluded.

### Proteomics data processing

The UNC Proteomics Core Facility processed raw data using Proteome Discoverer (Thermo Scientific, version 2.5). Data from the human samples were searched against a reviewed Uniprot human database (downloaded January 2022, containing 20,360 sequences), and data from the mouse samples were searched against a reviewed Uniprot mouse database (downloaded January 2021, containing 17,051 sequences) using the Sequest HT search algorithm within Proteome Discoverer. Enzyme specificity was set to trypsin, up to two missed cleavage sites were allowed, carbamidomethylation of Cys was set as a fixed modification and oxidation of Met was set as a variable modification. Label-free quantification (LFQ) using razor + unique peptides was enabled. Proteins were filtered for a 1%/5% peptide/protein level false discovery rate (FDR), and a minimum of 2 peptides. Further analyses were performed in Perseus (version 1.6.14.0), GraphPad, and DAVID bioinformatics. Gene Ontology Cellular Component was used to annotate EV-derived proteins. Only proteins with fewer than 50% missing values across samples were considered for statistical analysis. Missing values were imputed from a normal distribution with width of 0.3 and downshift of 1.8. Student’s t-test was performed for each pairwise comparison (mouse burn_control; human burn_control) and a p-value < 0.05 was considered statistically significant. A LFQ log2 fold change ratio for each pairwise comparison was calculated and a log2 ratio ±-1 was considered significant.

### Other Statistical Analysis

Analysis was conducted after data normality was met using D’Agostino & Pearson. Continuous variables were compared using Mann-Whitney or Kruskal-Wallis with Dunn’s multiple comparison. Correlation was performed with Spearman rank. Analysis was performed using GraphPad Prism v9.0 (La Jolla, CA).

For nanoString, negative binomial mixture, simplified negative binomial, or log-linear models were used depending on each gene’s mean expression compared to the background threshold. Multiple testing correction was performed using the method of Benjamini-Yekutieli. Causal Network Analysis was performed using IPA (Germantown, MD).

## RESULTS

### Plasma EVs from human burn injury patients promote cytokine responses in macrophages

Previously, we reported that burn EVs isolated from murine burn models can induce cytokine changes in various immune cell types (6) and reprogrammed immune responses to PAMPS such as LPS. We discovered that EVs isolated at different time points after injury have different immunomodulatory effects. We have built upon this to examine immunomodulation by human plasma EV isolated from human burn patients (burn-EVs) early (<72 after injury) and late (7-14 days after injury). We added equivalent numbers of EVs (3 x 10^7^) isolated from burn patients or healthy controls to human PMA-conditioned THP-1 macrophages (Figure 1A). After 48 h of culture, we measured media supernatant cytokines and chemokines *via* multiplex analysis. EVs from healthy subjects (“healthy-EVs”) and patients early (“early burn-EVs”) or late (“late burn-EVs”) after burn injury were able to promote differentially significant cytokine and chemokine secretion changes. As we have described before in mouse models, in the absence of LPS early– and late burn-EVs increased secretion of IL-6, IFNᵧ, TNF and CCL2 (MCP-1) compared to healthy-EV controls and cultures without EV addition (Figure 1A). IL-1β was uniquely induced at significantly higher concentrations by late burn-EV compared to early burn-EV and other controls. Other cytokines and chemokines evaluated were not significantly altered by EV exposure regardless of EV source. These data indicate that burn-EVs can alter macrophage immune responses, more specifically early injury burn-EVs promote a similar cytokine profile that is clinically observed after burn injury (2, 14).

**Figure 1:**
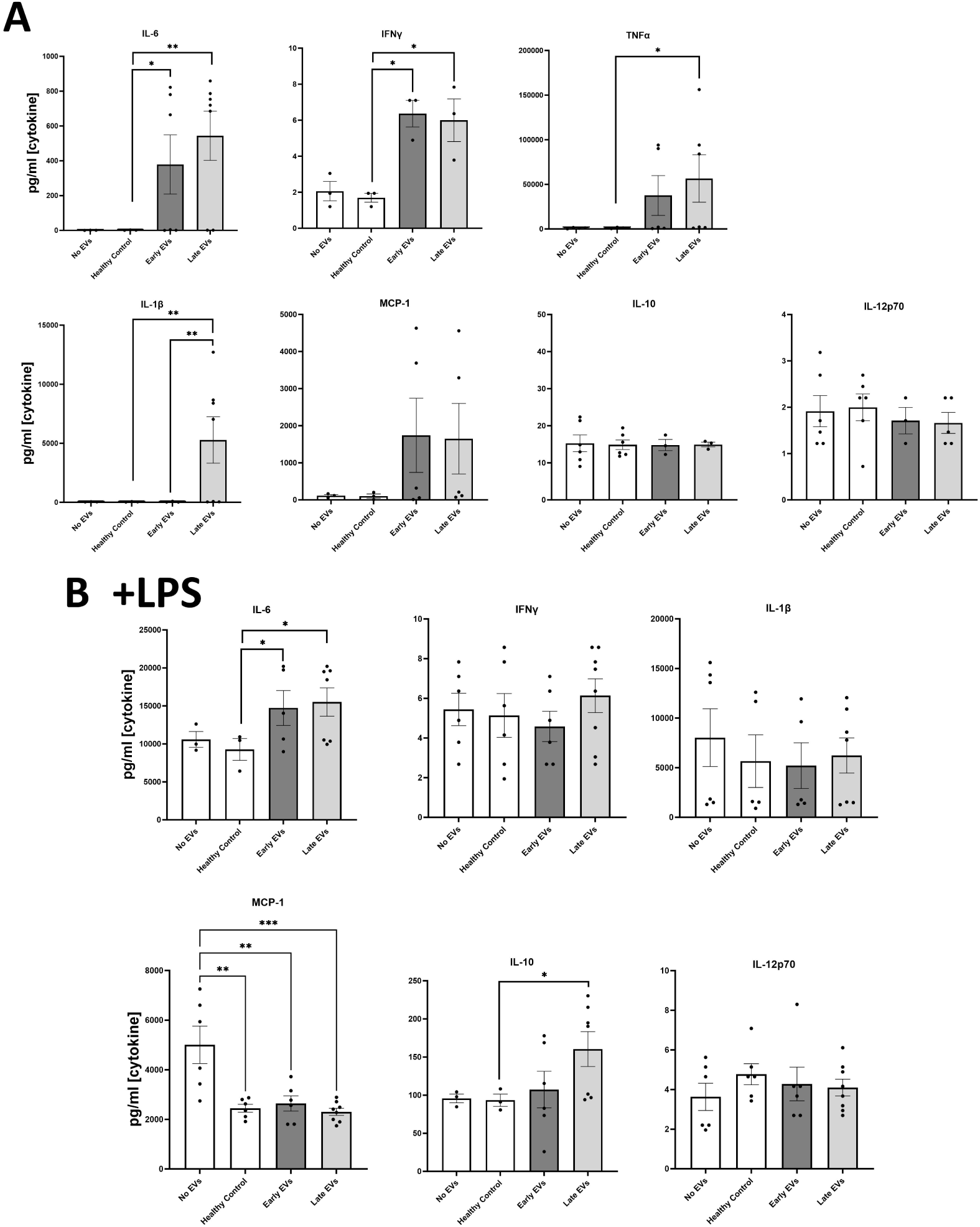
Human extracellular vesicles (EVs) isolated after burn injury regulate cytokine and chemokine release from PMA-conditioned THP-1 macrophages. EVs (3×10^7^ EVs) were isolated either “early”, 0-72 hrs, or “late” 14-20 days after injury or from healthy controls and applied to cultures for 48 hours (A) in the absence of or (B) presence of LPS to mimic burn-induced sepsis. Media cytokines and chemokines were measured by multiplex ELISA. Data shown as mean ± SEM; *\** p < 0.05, ** p<0.01, *** p<0.0001.

We have reported that EVs isolated late after burn injury can induce reduced phagocytic function in macrophages which could explain in part the increased susceptibility to infections in mice (15). Increased infection risk is associated with post-burn pathology, even acutely after injury (2), which can be mimicked by LPS addition. Therefore, we tested the impact of early and late human burn-EVs on THP-1 macrophages in the presence of LPS (Figure 1B). After 48 h of culture, we found that early and late burn-EVs induced significantly more secretion of IL-6 compared to healthy EV controls and no EV cultures (Figure 1B). CCL2 (MCP-1) secretion was significantly reduced by presence of EVs (healthy or burn-EV). However, the only secreted factor that was significantly altered by burn EVs uniquely was IL-10 which was significantly increased in the presence of EVs harvested late after injury compared to no EVs, healthy, or early burn EVs cultures. These data indicate that burn EVs can alter macrophage immune responses to LPS, with a significant shift towards an IL-10 / IL-6 phenotype, a similar cytokine profile that has been correlated with poor outcomes after burn injury (2, 14).

### Human early burn-EVs induce proinflammatory reprogramming of THP-1 macrophages

The clinical phases after burn injury are defined as an early pro-inflammatory “shock” phase and a later compensatory phase associated with significant susceptibility to infection. The mechanisms underlying these phases are not as clearly defined immunologically. One characteristic of burn injury is significant immune dysfunction, with macrophages being impacted greatly, resulting in more susceptibility to infection and mortality. To determine the impact of early burn-EVs on macrophage activation, plasma EVs isolated from patient collected within 3 days after burn injury or healthy controls (3×10^7^) were added to PMA-conditioned THP-1 macrophage cultures (5×10^6^ cells/well). After 48 hours of culture, RNA was purified from cell lysates for nanoString multiplex immune gene expression analysis. Early human burn-EVs caused a significant upregulation of many immune genes in THP-1 cells compared to healthy control EVs. To elucidate differences in immune signaling pathways affected by early burn-EVs versus healthy controls, we performed Ingenuity Pathway Analysis (IPA). Using predicted Z-scores, early burn-EVs significantly altered the activity of several pathways compared to the healthy EVs (Figure 2B). This included increased activation in IL-6 Signaling, T_h1_, Acute Phase Response Signaling and Macrophage Classical Activation Signaling Pathways (p <0.05). Decreases in IL-10 and Macrophage-Stimulating Protein (MSP)/ Recepteur d’Origine Nantais (RON; MSP-RON) Signaling in Macrophage Pathways were also observed. This is consistent with early burn-EVs inducing pro-inflammatory signals in macrophages compared to healthy EVs. We then tested the impact of early burn-EVs on THP-1 activation in the presence of LPS using nanoString mRNA analysis. Human early burn-EVs caused less robust transcriptional response beyond the LPS effect (Figure 2C). In total, six genes were significantly altered by burn-EVs in the presence of LPS (3 genes upregulated, 3 genes downregulated). Together, this indicates that early burn-EVs induce pro-inflammatory signaling in human THP1 macrophages that is diminished in the presence of LPS.

**Figure 2.**
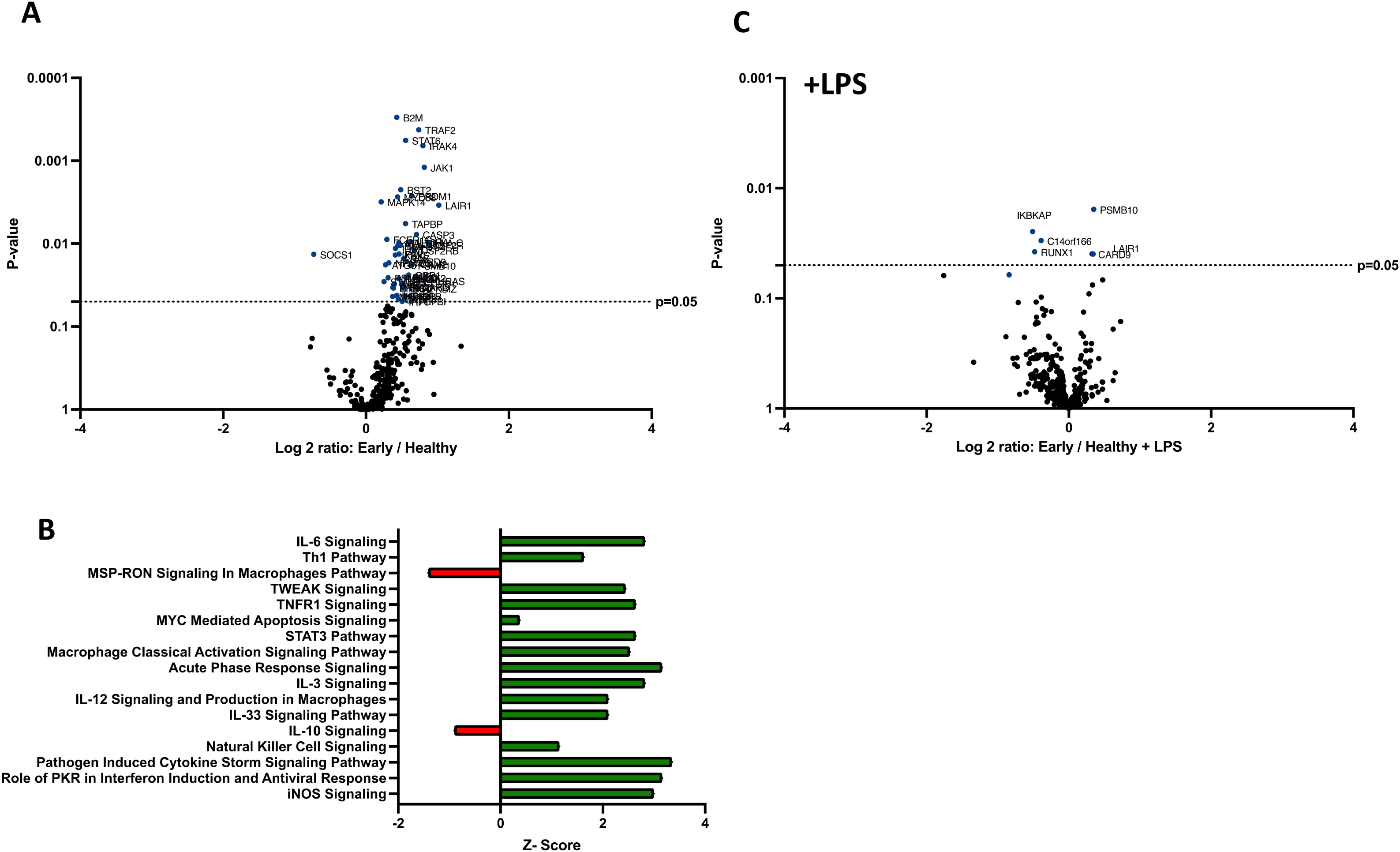
Human extracellular vesicles (EVs) isolated early after burn injury reprogram THP-1 macrophages to a proinflammatory state. PMA-conditioned THP-1 cells were exposed to 3×10^7^ EVs /well from human burn patients (<72 hrs after injury) or healthy controls in the absence (A-B) or presence (C) of LPS. Gene expression was evaluated using nanoString barcoding spanning 561 mRNAs (nCounter Human Immunology CodeSet v2.0). Data are presented as the log2-transformed differential fold change in immune gene expression (A and C), with associated p-value significance (using Welch’s t test), after data normalization to housekeeping and internal control genes by nSolver v4.0. Differential fold change is shown as early EV *versus* healthy EVs (each group represents cell cultures stimulated with EVs from 6 patients). Differential gene expression and pathway Z-scores between early burn-induced EVs and healthy controls were analyzed via Ingenuity Pathway Analysis (B); only significantly altered (p<0.05) pathways are shown.

### Human late burn-EVs induce anti-inflammatory reprogramming of THP-1 macrophages

In previous studies, we have described a compensatory phase (day 14) after a 20% TBSA burn in mice (7). To determine if late burn-EVs elicit anti-inflammatory responses in macrophages, we probed mRNA from PMA-conditioned THP-1 macrophages stimulated for 48 hours in the presence of 3 x 10^7^ plasma EVs isolated 18-25 days after burn injury in human patients or from healthy controls. To understand what genes and pathways were being affected by late burn-EVs, we again turned to nanoString analysis. Late burn-EVs caused a significant downregulation in the expression of several immune genes in THP-1 macrophages (Figure 3A; 1 upregulated, 28 downregulated). IPA found that several pro-inflammatory pathways were downregulated with a corresponding upregulation of anti-inflammatory IL-10 signaling pathways. Similar findings were observed even in the presence of LPS, with late burn-EVs causing a significant down regulation of many immune genes in the presence of LPS compared to EV from healthy donors (Figure 3C, 0 upregulated, 60 downregulated). IPA revealed significant downregulation of multiple LPS-induced signaling pathways by late burn-EVs (Pathogen Induced Cytokine Storm, T_h1_ and T_h2_, NFkB signaling pathway), with upregulation of anti-inflammatory IL-10, PPAR and PTEN signaling (Figure 3D).

**Figure 3.**
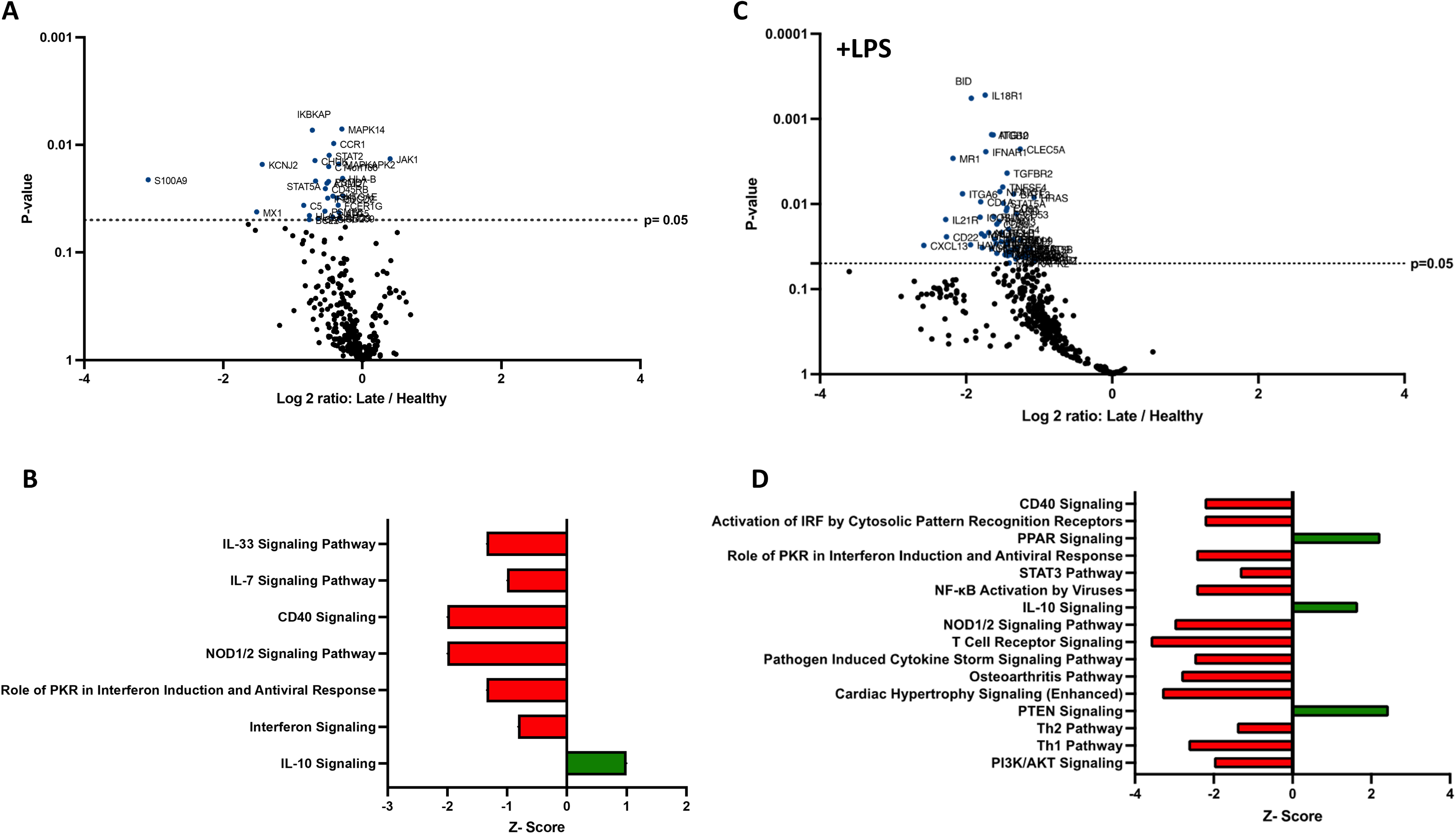
Human extracellular vesicles (EVs) isolated late after burn injury reprogram THP-1 macrophages to an anti-inflammatory state. PMA-conditioned THP-1 cells were exposed to 3×10^7^ EVs/well from human burn patients (14-20 days after injury) or healthy controls in the absence (A-B) or presence (C-D) of LPS. Gene expression was evaluated using nanoString and are presented as the log2-transformed differential fold change in immune gene expression (A and C), with associated p-value significance (using Welch’s t test), after data normalization to housekeeping and internal control genes by nSolver v4.0. Differential fold change shown early EV *versus* healthy EVs (each group represents cell cultures stimulated with EVs from 6 source patients, with n=6 different EV preparations from 2 experiments). Differential gene expression and pathway Z-scores between early burn-induced EVs and healthy controls were analyzed via Ingenuity Pathway Analysis (B and D); only significantly altered (p<0.05) pathways are shown.

Compared to early burn-EVs, late burn-EVs caused a profound downregulation of immune genes both in the absence (Figure 4A, 0 upregulated, 152 downregulated) and presence of LPS (Figure 4C, 0 upregulated, 110 downregulated). Significant pathway changes including reduction in Macrophage Classical Activation Signaling Pathway, NK cell signaling, iNOS signaling, with a significant increase in IL-10 signaling by late burn-EVs versus early burn-EVs (Figure 4B, D).

**Figure 4.**
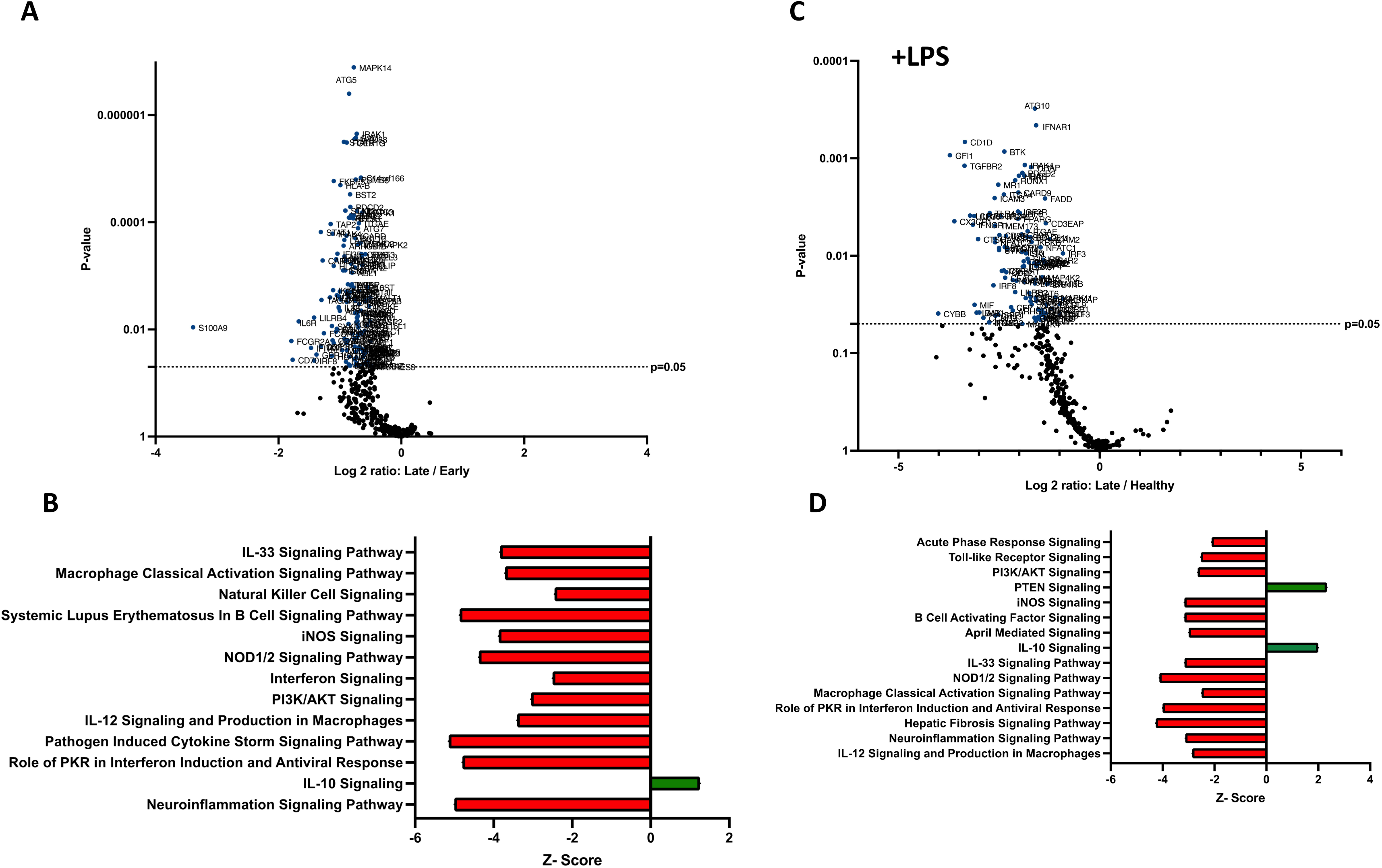
Human extracellular vesicles (EVs) isolated early and late after burn injury induce differential reprogramming in THP-1 macrophages. Gene changes induced in THP-1 cells by EVs from human burn patients isolated late (14-20 days) compared to human burn patients isolated early (0-3 days), in the absence (A-B) or presence (C-D) of LPS. Differential fold change is shown as late *versus* early EVs (each group represents cell cultures stimulated with EVs from 6 source patients, with n=6 different EV preparations from 2 experiments). Differential gene expression and pathway Z-scores between early burn-induced EVs and healthy controls were analyzed via Ingenuity Pathway Analysis (B and D); only significantly altered (p<0.05) pathways are shown.

### Mouse burn-EV cargo changes temporally in a manner consistent with clinical course

We have previously reported that the burn EV protein content changes early after burn injury in mice and in humans (0-3 days after injury) compared to healthy controls (6, 15). However, the changes in protein content at later time points after injury are not known. In this follow upstudy, we aimed to characterize the proteomes of EVs purified late after burn injury in humans and mice. Bioinformatics analysis was also performed to identify putative signaling pathways impacted by the EV protein cargo early and late after burn injury in both humans and mice. LC-MS/MS measured 1080 proteins in the EV samples. Seven days after injury the protein cargo of burn-EVs was significantly altered (Figure 5A, 100 proteins upregulated, 37 proteins decreased). At day 7, the majority of the upregulated proteins were membrane, circulation/complement, neuronal and protein-regulating proteins, while the majority of the downregulated proteins were membrane-associated (Figure 5B-C). At day 14 after injury, however, only 24 proteins were increased, while 152 proteins were decreased in burn EVs (Figure 6A). A few proteins were increased in similar categories as at day 7, however a vast down regulation of membrane, metabolic, protein regulation, and circulation/complement proteins was seen. Further, a broad downregulation in proinflammatory pathways was predicted by IPA.

**Figure 5.**
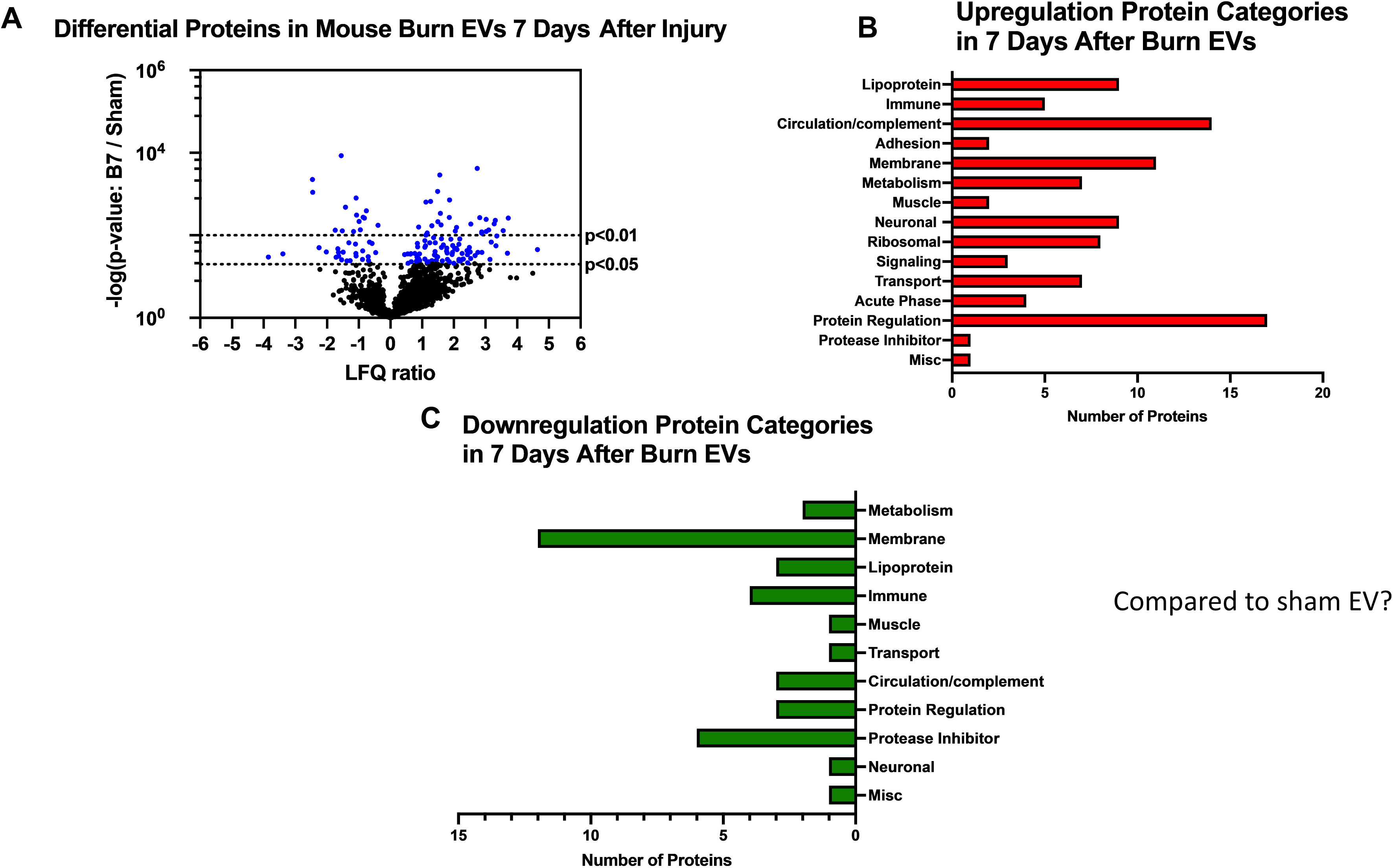
Burn injury in mice alters protein cargo of plasma EVs 7 days after injury. Adult mice underwent a 20% total body surface area (TBSA) burn and plasma EVs were collected 7 days after injury and protein content measured by LC-MS/MS. (A) Differentially expressed protein peptides in burn *versus* sham EVs (LFQ ratio, p-value Student’s t-test, N=3 per group). (B) Characteristics of proteins that are increased in burn EVs relative to controls. (C) Characteristics of proteins that are decreased in burn EVs relative to sham controls.

**Figure 6.**
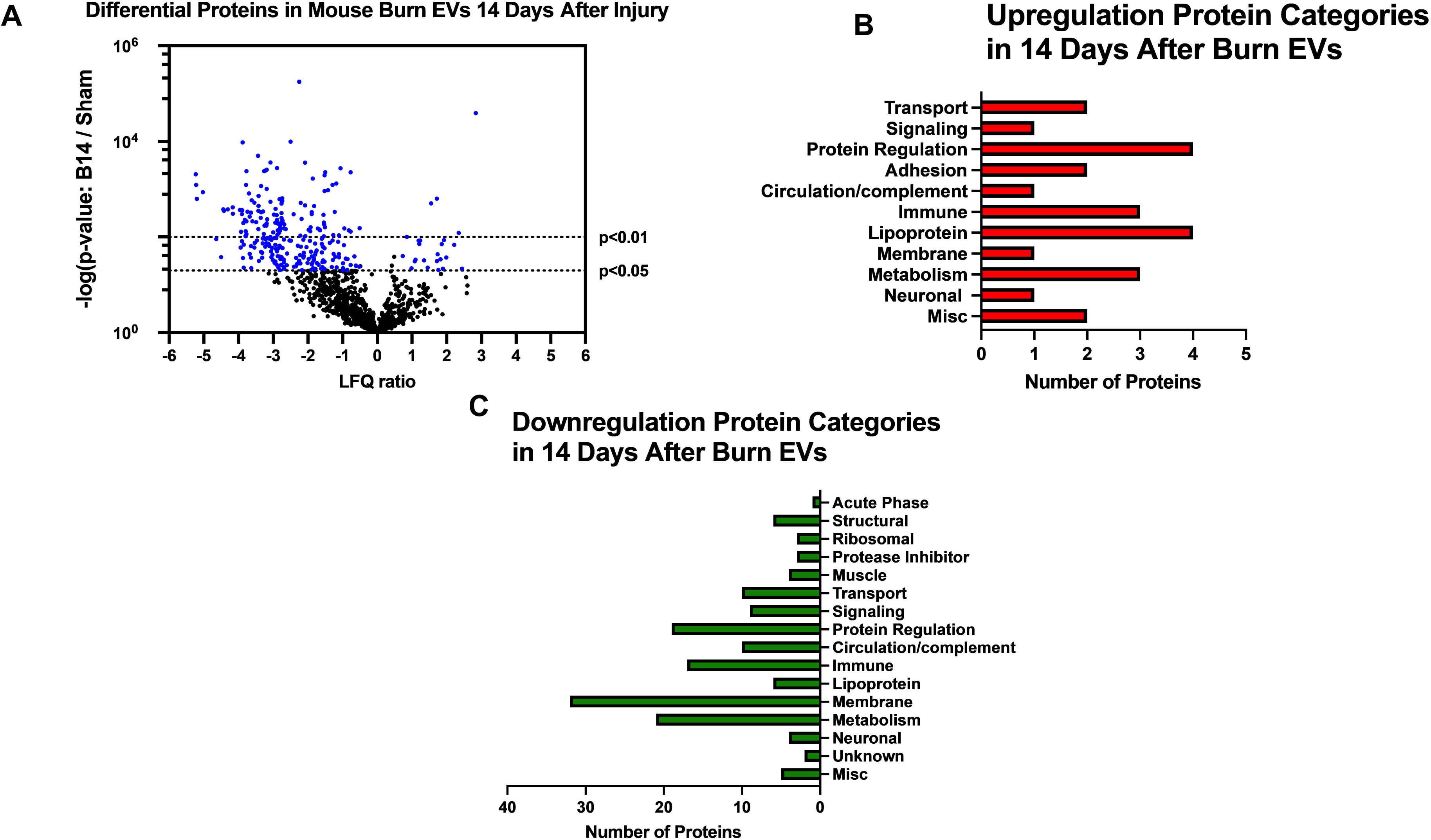
Burn injury in mice alters protein cargo of plasma EVs 14 days after injury. Adult mice underwent a 20% total body surface area (TBSA) burn and plasma EVs were collected 7 days after injury and protein content measured by LC-MS/MS. (A) Differentially expressed protein peptides in burn *versus* sham EVs (LFQ ratio, p-value Student’s t-test, N=3 per group). (B) Characteristics of proteins that are increased in burn EVs relative to controls. (C) Characteristics of proteins that are decreased in burn EVs relative to sham controls.

—Utilizing these day 7 and day 14 datasets, as well as the previously collected dataset from day 3 (6), we performed IPA to identify the putative pathways impacted across the three time points after injury compared to sham mice (Figure 7). Using IPA, proteins from burn-EVs from day 3 were predicted to increase coagulation and intrinsic prothrombin activation pathways (Figure 7A). At day 7 after injury, more pathways were upregulated, including post-translational protein phosphorylation, coagulation, LXR/RXR activation, APR signaling, production of nitric oxide and reactive oxygen species, TLR signaling, activation of phagocytes, and pathogen induced cytokine storm signaling pathways (Figure 7B). Day 14 burn-EVs were predicted to have broad downregulation of numerous pathways (26 pathways significantly downregulated, Figure 7C). This includes phagosome formation, mTOR activation, and Fcγ receptor-mediated phagocytosis by macrophages and monocytes. RHOGDI signaling was the only pathway predicted to be upregulated.

**Figure 7.**
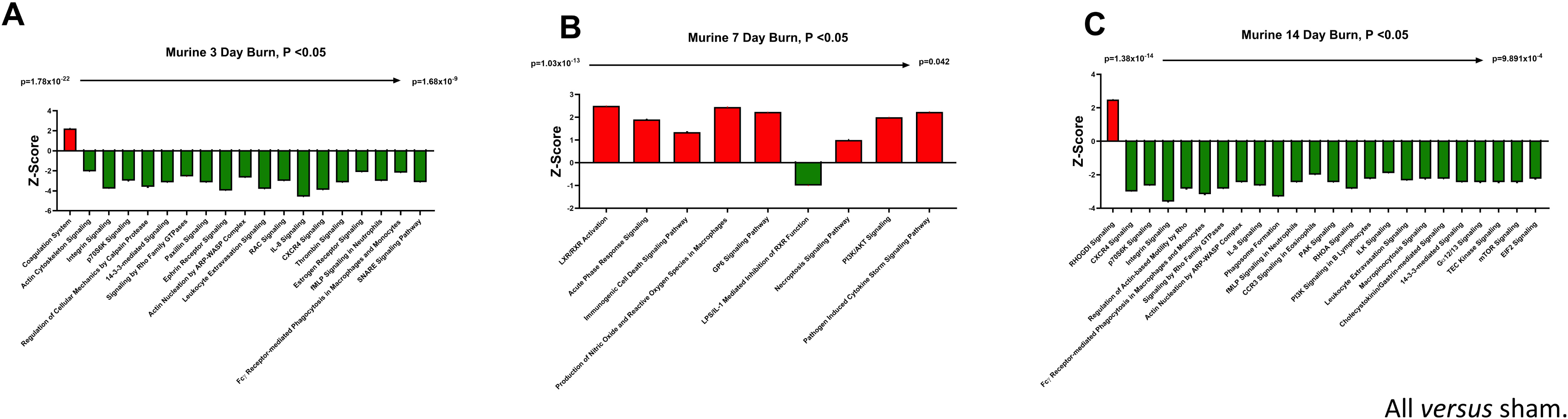
EV protein cargo pathways altered over time after burn injury in mice. Adult mice underwent a 20% total body surface area (TBSA) burn and plasma EVs were collected at different time points after injury and protein content measured by LC-MS/MS (p-value Student’s t-test, N=3 per group). Differential protein expression and pathway Z-scores between burn at different time points past injury; (A) 3 days following injury (data reanalyzed from Maile *et al* 2021(6)), (B) 7 days following injury and (C) 14 days following injury and sham were analyzed via Ingenuity Pathway Analysis; only significantly altered (p < 0.05) pathways are shown.

We next identified specific proteins that either changed or were stable over time after burn injury. Levels of eight common proteins significantly altered at 3 and 7 days after injury and 22 protein levels were significantly altered at both 7 and 14 days post-injury (Figure 8A). Levels of five common proteins were significantly altered at both 3 and 14 days after injury, and only one protein at all data points. However, the directionality of change of these 36 proteins varied at each time point (Figure 8B). For instance, several Ig kappa chains showed a progressive reduction in their levels in EVs from 3 to 14 days, while several apoliproteins were not detected at 3 days but increased at both 7 and 14 days after burn. This indicates that in the EV compartment circulating levels of several apolipoproteins are increased, while production of IgGs is reduced. EVs could represent a reservoir or sponge for IgGs and apolipoproteins. Pathway analysis showed that certain predicted pathways followed a U-shaped curve (PI3K/AKT signaling, phagosome formation and GP6 signaling) while LXR/RXR showed a persistent reduction (Figure 8C). Interestingly, the pattern of Phagosome Formation is similar to clinical observations, wherein the “mid” 7 day point is often regarded as a inflection point between the acute shock and PICS phase in mouse models of burn injury (1). Others followed an inverted U shaped pattern (induction of iNOS and ROS in macrophages) while Pathogen induced Cytokine Storm signaling and Intrinsic Prothrombin Activation were high early and returned to baseline at 14 days.

**Figure 8.**
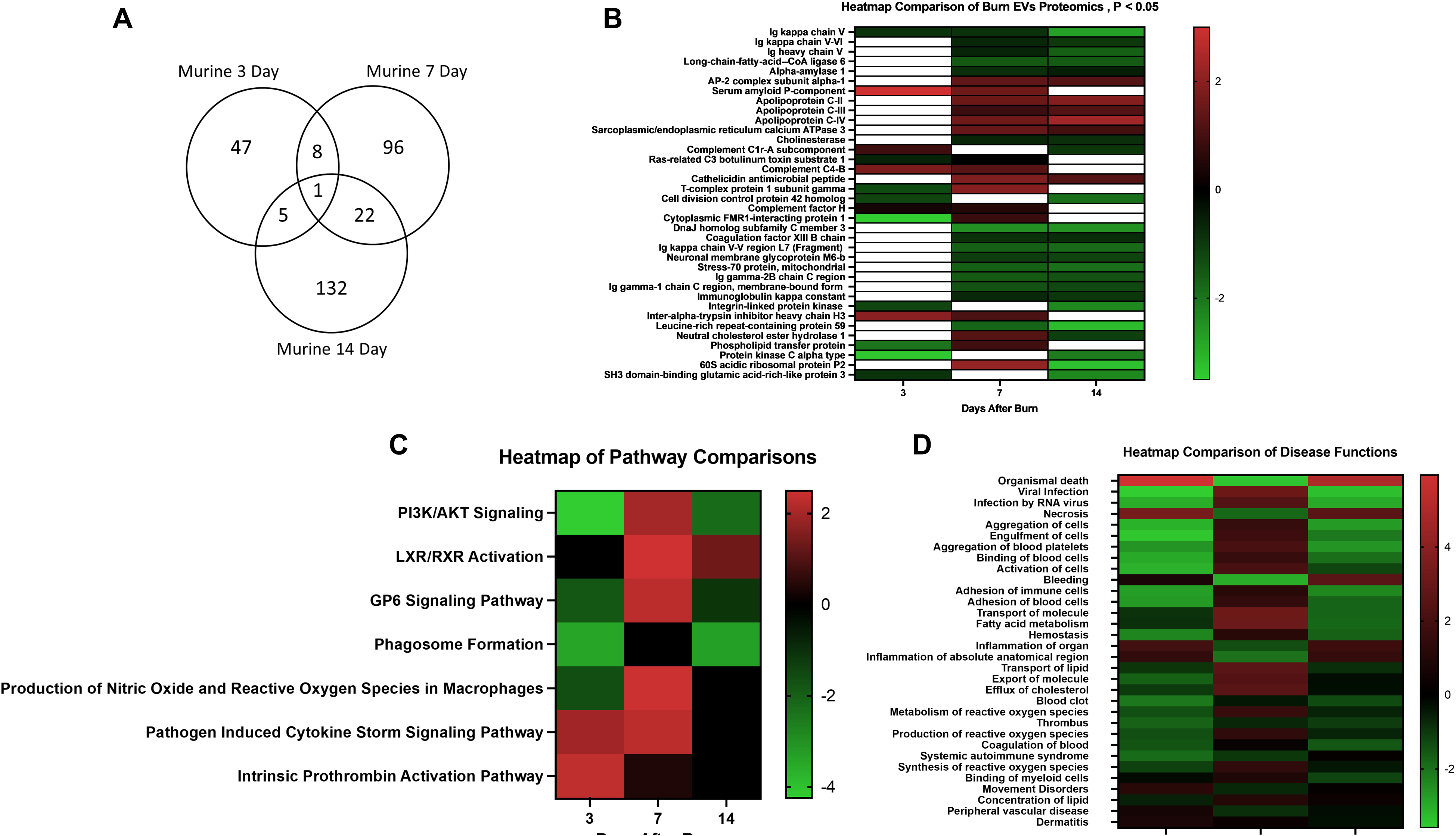
Comparison of significantly expressed proteins in mouse plasma EVs over time after burn injury. Adult mice underwent a 20% total body surface area (TBSA) burn and plasma EVs were collected at early (3 days), intermediate (7 days) and late (14 days) time points after injury. Protein content was measured by LC-MS/MS. A Venn diagram in (A) summarizes the number and identity of proteins between the different time points after injury (p-value Student’s t-test, N=3 per group). (B) Heatmap showing Z-scores of protein expression between burn and sham, red shows positive Z-score > 1, and green shows negative Z-score < –1. (C) Heatmap showing Z-scores of pathways from EV proteome between burn and sham, red shows positive Z-score > 1, and green shows negative Z-score < –1. (D) Heatmap showing Z-scores of disease function from EV proteome between burn and sham, red shows positive Z-score > 1, and green shows negative Z-score < –1. White denotes no significant fold change detected.

Predicted disease functions also followed various patterns, with predicted organismal death being high early and late but low at 7 days, similar to clinical natural history in human burn patients (Figure 8D).

### Temporal changes in human burn EVs provide insight into clinical progression after injury

We next compared EV cargo in human burn patients isolated early (<3 days) and late (>14 days) after injury. To control for patient to patient variability, late plasma samples for each patient were compared to early samples for that same patient. Overall, 446 proteins were measured in human EVs. Of these, 183 were significantly altered in late burn-EV versus early burn-EV, with 172 increased and 11 decreased (Figure 9A). Proteins that increased from early to late after injury were mainly immune proteins, circulation/complement, and transport proteins (Figure 9B), while those with reduced levels were mostly lipoproteins (Figure 9C). Pathway analysis on our previously published early burn-EVs (compared to healthy-EV) predicted several reductions including LXR/RXR activation (Figure 9D), which was predicted to be reduced in early mouse EVs as well (Figure 8C). Also similar to mouse burn-EVs isolated early (3 days) after burn injury, PI3/AKT signaling was predicted to be increased which was also seen in mice. At later time points after injury, LXR/RXR activation was increased similar to mice. Further, Leukocyte Extravasation, Phagosome Formation, and IL-8 signaling were all increased, though Production of iNOS and ROS in Macrophages was predicted to be reduced (Figure 9E). RHOGDI signaling, which is critical for leukocyte migration was reduced. Upstream analysis predicted activation of TGFβ signaling (z score: 3.7, *p* = 7.7 x 10^-46^) and IL-1β (z score: 3.4, *p* = 1.97 x 10^-24^) and an inhibition of PSEN1 (z score: –2.1, *p*=5.16 x 10^-25^).

**Figure 9.**
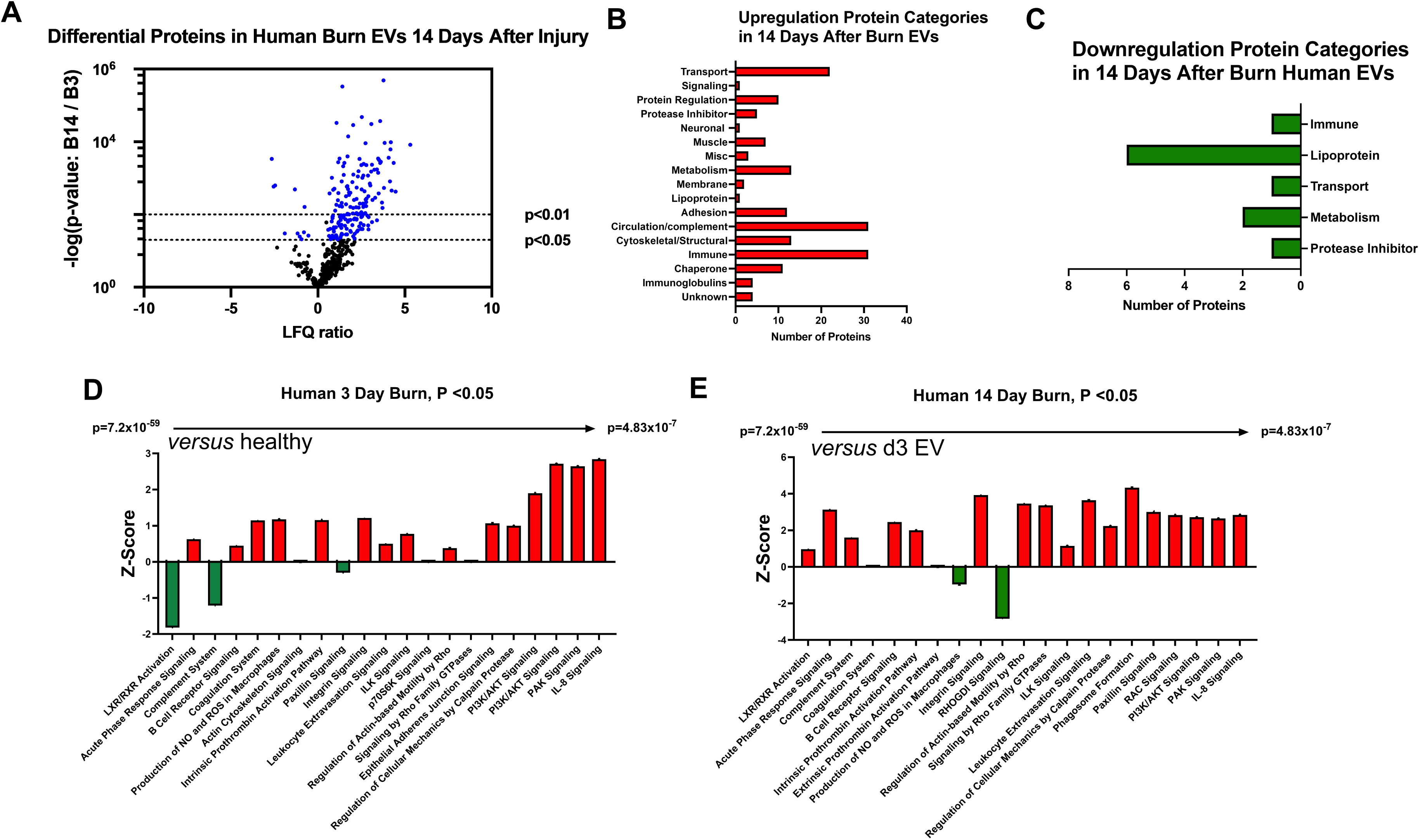
Burn injury alters protein cargo of human plasma EVs 14 days after injury. Plasma EVs from human burn patients with severe burn injury isolated 14-20 or 3 days after admission. Protein content was measured by LC-MS/MS. (A) Differentially expressed protein peptides in late after burn (14-20 days) vs 3 days following admission (LFQ ratio, p-value Student’s t-test, N=3 per group). (B) Characteristics of proteins that are increased in burn EVs 14-20 days relative to 3 days. (C) Characteristics of proteins that are decreased in burn EVs relative to controls. Ingenuity Pathway Analysis quantified pathway Z-scores between burn at different time points past injury; (D) 3 days following injury compared to healthy, (E) 14 days following injury compared to 3 days following injury; only significantly altered (p < 0.5, n=3 per group) pathways are shown.

## DISCUSSION

Burn injury results in profound immune, metabolic, and coagulopathic dysregulation with chronic persistent immune and physiological derangements. After resolution of the initial burn shock, morbidity and mortality occurs weeks after injury mostly due to infectious complications (16–23). Gaps in the knowledge that would directly improve patient care include identification of molecular mediators that could be targeted for therapeutic intervention to normalize immune function and physiology, and discovery of markers that would identify high risk individuals with immune dysfunction at the time of injury.

In this study we completed a thorough analysis of plasma EVs that are released after burn injury that we have demonstrated here and previously (6–8) promote immune dysfunction and can be potentially be used as biomarkers to identify at-risk patients. Using both a mouse model of burn injury and patient samples we find that plasma EVs are altered in their cargo and immunoregulatory activity after burn injury. These EVs contain immunomodulatory molecules such as key proteins lipids, DAMPS, miRNAs and cytokines. We have identified clear immunomodulatory effects on immune cells by EVs harvested early (<3 days) and late after injury (14 days) in our 20% TBSA mouse burn model (6). In both systems, EV protein cargo was assessed in an unbiased manner by mass spectrometry, and identified cargo overlap between humans and mice that correlated with eventual length of hospital stay (6). We also found that transfer of early burn-EVs to uninjured mice reproduces a similar systemic immune response that is seen with burn injury itself (7). Here, we found that human EVs purified early and late after burn injury reprogram immune responses in human macrophages in a manner consistent with the known clinical course with burn injury. Early burn-EVs after burn injury in humans and mice promote proinflammatory signaling and are distinct in their cargo from late burn-EVs which have anti-inflammatory activity. We hope that these and future efforts in larger numbers of patients will be able to identify specific molecular EV fingerprints that can identify high risk patients and perhaps guide treatment management. Using detailed bioinformatic analysis of EV protein cargo across the timeline after injury, we characterized the putative immune and physiologic pathways that burn-induced EV impact. Taken together, these data suggest burn-EVs might be useful in biomarker and therapeutic development.

These observations are in agreement with O’Dea et al. who have found that elevated levels of leukocyte– and granulocyte-derived EVs correlate with clinical assessment scores of burn severity associated with the risk of severe sepsis (24). In addition to immune responses, burn-induced EVs have been found to promote endothelial barrier dysfunction which could promote post-burn pathology in the lungs and gut (25). We and others have also demonstrated that macrophages play a key role in both early and late immune dysfunction (14, 26–36) with aberrant macrophage cytokine and chemokine function leading to tissue damage driving further burn-induced immune dysfunction. Conversely, aberrant phagocytosis by macrophages (which we found occurs in response to mouse late burn-EVs) can promote increased susceptibility of infection, a devastating consequence of immune dysfunction after burn-injury (33, 34, 37). Using unbiased immune transcriptome analysis and causal network bioinformatic analysis, we found that early human burn-EVs induce reprogramming in human macrophages *in vitro*, with a very different immunogenic profile to that observed with late burn-EVs. Of particular importance was the significantly increased activation in IL-6 Signaling. In burn patients, sustained high levels of circulating IL-6 corelate strongly with poor outcomes after burn injury (38–40). These are all hallmarks of the observed SIRS phase after burn injury. There was also significantly decreased activation of IL-10 and MSP-RON Signaling in Macrophage Pathways, in accordance with our findings in mouse models (28). In contrast, EVs isolated late after burn injury induced a large scale downregulation of pro-inflammatory pathways and a corresponding upregulation of anti-inflammatory IL-10 signaling pathways. These data support our findings, in the burn mouse model, that EVs from burn-injury induce a unique set of immune genes in transformed and primary macrophages compared to EVs from sham-injured mice and that burn-EVs isolated at distinct phases of burn-injury have differential immunomodulatory effects.

We also found that late human burn-EVs significantly altered macrophage responses to LPS *in vitro* compared to EVs from healthy subjects, with the most marked effect being the significant downregulation of multiple LPS-induced signaling pathways by late burn-EVs (Pathogen Induced Cytokine Storm, NFkB signaling), and upregulation of anti-inflammatory IL-10, Peroxisome proliferator-activated receptor (PPAR) and PTEN signaling. This could contribute to the immunosuppressive phase seen clinically. Conventional dogma suggests that molecular feedback inhibition underlies the transition from proinflammatory to anti-inflammatory signaling. However, this work suggests that changes in EV cargo and activity may also contribute. For instance, we previously reported that early after burn tissues release danger signals that induce acute activation of several signaling pathways including Mammalian Target of Rapamycin (mTOR) and NFκB (6, 8, 14, 15, 26, 41–43) which drive the execution of metabolic cellular programing and inflammatory functions causing further local and systemic damage, and SIRS (44). mTOR activation leads to the rapid upregulation of the negative regulator of inflammation, PPARγ, which in turn represses mTOR (45), while also promoting anti-inflammatory pathways (44, 46–48). Similarly, NFκB-mediated transcription induces the upregulation of its negative regulator, IκB, and PPARγ also regulates the NFκB pathway through both a physical interaction preventing NFkB nuclear translocation, and active transcription of IκB (49) which we have also recently shown occurs in circulating immune cells in patients with severe burns (7). Here we present new data that human burn-EVs from burn patients do indeed also induce a unique set of immune genes in macrophages compared to EVs from healthy individuals and that burn-EVs isolated at distinct phases of burn-injury have differential immunomodulatory effects, both of which are reminiscent of the temporal immunological phenotypes observed clinically (2, 8, 14, 26–29, 31, 32, 41, 43). Together these data implicate EV-macrophage interactions in the burn-induced immune dysfunction.

Using unbiased mass spectrometry techniques and bioinformatic causal pathway analysis, we found that human and mouse burn-EVs isolated at different timepoints after burn injury have unique and overlapping protein burn-EV cargo and predicted impacted pathways. Our published data demonstrated that there are some commonalities between the protein components in both human and mouse burn-EVs purified early after burn injury, such as the acute phase response protein SAA1 and increases in circulating complement/coagulation factors (6). Here, we found at day 7, burn-EVs were enriched in circulation/complement, neuronal and protein-regulating proteins. We are currently exploring the role of neuronal cell markers as an indicator of cell origin of EV that arise after burn injury. At day 14, the upregulated proteins had a similar pattern as day 7, but the upregulated proteins the majority were metabolism-associated proteins. These patterns of protein cargo predicted specific pathways that were impacted compared to EVs from sham injured mice. We observed a shift in burn-EVs from day 3 (mostly downregulated pathways, with upregulation of coagulation, intrinsic prothrombin activation pathway, and pathogen induced cytokine storm), to day 7 (more upregulated pathways with increased LXR/RXR activation, PI3K/AKT, and production of iNOS and ROS), to day 14 (mostly downregulated pathways with only RHOGDI signaling significantly upregulated). Analysis of early human burn-EVs predicted upregulated of several clinically relevant pathways (coagulation, IL-8, PAK and production of iNOS/ROS) with downregulation of complement and LXR/RXR activation. Late burn-EVs rather were predicted to have upregulated complement and LXR/RXR activation, with decreased in RHOGDI and production of iNOS/ROS. There are clear differences in the directionality of pathways that are potentially impacted between mice and humans, but it is well described that the kinetics of immune dysfunction differ between humans and mice after burn injury(35). These data, however do implicate that these pathways are of importance in the post-injury response in both humans and mice, which deserves greater research attention.

Liver X receptors (LXR) and retinoid X receptors (RXR) are a receptor family of transcription factors closely related to nuclear receptors such as PPAR. LXRa is expressed in macrophages and forms heterodimers with the obligate partner retinoid X receptor and can be activated by lipid ligands such as oxysterol derivatives of cholesterol to regulate inflammatory and metabolic responses. This pathway has been implicated in mediating coagulopathy in trauma (50), and sex differences in other disease systems but no study has investigated the role of LXR/RXR signaling after burn injury. RHOGDI (Rho GDP-dissociation inhibitor) are negative regulators of RHO GTPases, preventing their activation. RHO GTPases have been implicated in the regulation of endothelial permeability and leukocyte migration through actin cytoskeletal organization (51, 52). Oxidative stress, as seen after burn injury and trauma, regulate barrier function via RHO GTPases (53). This study has therefore implicated these molecular regulatory compounds as potential modes of action of EVs that arise after burn injury and require further investigation.

We, and others, have also reported that in human and pre-clinical studies macrophages play a key role in both early and late immune dysfunction (14, 26–36) with aberrant macrophage cytokine and chemokine function leading to tissue damage associated with early burn-induced immune dysfunction. Conversely, aberrant phagocytosis by macrophages, can lead to susceptibility of infection; the hallmark of immune dysfunction observed during later stages of burn-injury (33, 34, 37). Indeed, we have published that burn injury results in a steady accumulation in the periphery of macrophages, which early after burn injury upregulate the innate immune receptors toll-like receptor (TLR) 2 and TLR4, followed by a decrease of TLR2 and TLR4 expression late after burn injury which resulted in temporal differences in TLR-induced cytokine responses (41). Here we present data that burn-EVs from burn-injury induce a unique set of immune genes in macrophages compared to healthy-EVs from sham-injured mice. Burn-EVs isolated at distinct phases of burn-injury have differential immunomodulatory effects, both of which are reminiscent of the temporal immunological phenotypes observed in our murine models of burn-injury and clinically (2, 8, 14, 26–29, 31, 32, 41–43). Together these data implicate EV-macrophage interactions in the burn-induced innate immune dysfunction.

Therefore, this and other work implicate EVs as critical mediators of immune activation after trauma. Studies assessing the further effects of human post-burn EVs on human immune cell function are needed, and we plan to do this using induced pluripotent stem cells (IPSCs) differentiated into human monocytes, neutrophils, epithelial cells, and endothelial cells. Together, findings presented here as well as recent work by others and ourselves indicate that EVs may be key and underexplored regulatory mediators in settings of trauma providing possible insight into cellular dysfunction. EV and associated cargo may be used to identify novel therapeutic targets.

## Scope

This manuscript highlight the ability of extracellular vesicle (EV) cargo to alter significantly over time after trauma, which alters their ability to immunomodulate innate immune cells. We performed this study after burn injury in both mouse models and human patients. This is of general interest to the immune field, as EVs have emerged as key drivers of inflammation in many systems.

## Abbreviations

AAALAC: American Association for Accreditation of Laboratory Animal Care
CARS: Compensatory Anti-Inflammatory Response Syndrome
DAMPs: Damage-Associated Molecular Patterns
EVs: Extracellular vesicles
HMGB1: High-mobility group protein 1
HA: Hyaluronic acid
LPS: Lipopolysaccharide
MARS: Mixed Agonist Response Syndrome
NTA: Nanoparticle Tracking Analysis
PICS: Persistent Inflammation, Immunosuppression, and Catabolism Syndrome
PRR: Pattern Recognition Receptors
SIRS: Systemic Inflammatory Response Syndrome,
TLR: Toll-Like Receptors
TBSA: Total Body Surface Area

## Acknowledgments

We thank the NC Jaycee Burn Center patients and families for their contribution.

